# Do These Crayfish Make Me Look Fat? Body Condition Correlates To Prey Abundance In Three Hellbender (*Cryptobranchus Alleganiensis*) Populations

**DOI:** 10.1101/659441

**Authors:** Kirsten Hecht, Max Nickerson, Michael Freake, Phil Colclough, Katie Stofer

## Abstract

In ecological studies body condition is often measured as an indicator of animal health or well-being. The Hellbender (*Cryptobranchus alleganiensis*) is a threatened salamander species found throughout the montane regions of the eastern United States. Although few young individuals have historically been found in the wild, recent studies in Blue Ridge Physiographic Regions have uncovered larvae in several streams. In Little River, Tennessee, differences in the crayfish population, the principal component of the adult Hellbender diet, was hypothesized as a potential reason for the large number of immature individuals, and lack of large adults. To investigate this hypothesis, we compared body condition of Hellbenders in three streams with different crayfish relative frequencies. Body condition of Hellbenders was positively correlated with crayfish relative frequencies, with Hellbenders in the stream with the highest crayfish relative frequency exhibiting the highest expected mass per total length.

---

Body condition metrics have traditionally been used in ecological studies to estimate animal health or ‘well-being’ (Anderson and Gutreuter, 1983; Stevenson and Woods, 2006). Body condition is generally defined as the deviation of an individual’s observed mass from the expected mass of an individual of the same length (Anderson and Gutreuter, 1983). Body condition is assumed to correlate with fat storage in the body (Pope and Matthews, 2002) and can correlate to health, fecundity (Girish and Saidapur, 2000), growth, and mortality (Anderson and Gutreuter, 1983). Although traditional molecular techniques can give a more accurate picture of body composition and health, these techniques often require invasive procedures, sacrifice of the study organism and/or removal from the wild, which may be undesirable in threatened and endangered species. These techniques can also be costly. Therefore, nondestructive estimators of body condition that can be easily measured in the field are still common in conservation biology (Stevenson and Woods, 2006).

Body condition is used as a tool to investigate ecological and conservation questions regarding amphibians (Stevenson and Woods, 2006) including the impacts of forestry practices on Plethodontid salamanders (Welsh et al., 2008; Homyack et al., 2011) and the effects of pesticides on anurans (Gendron et al., 2003; Levey, 2003). A number of factors can influence the body condition of amphibians including land use (Karraker and Welsh, 2006), stress, disease and parasites (Bodinof et al., 2012), food availability (Pope and Matthews, 2002), gender (Taber et al., 1975; Humphries and Pauley, 2005) temperature, and population density (Reading and Clark, 1995). Body condition appears to affect a variety of amphibian characteristics such as coloration (Davis and Maerz, 2007), breeding success (Howard et al., 1997), male calling (Howard and Young, 1998), reproduction (Lowe et al., 2006) and population dynamics (Welsh et al., 2008).

*Cryptobranchus alleganiensis*, a large (up to 740 mm) aquatic salamander, resides primarily in cool, oxygen-rich streams in the eastern United States (Nickerson and Mays, 1973a). The species is declining in many portions of its range. While large adults comprised the majority of captured Hellbenders in most study sites (Nickerson and Mays, 1973b; Taber et al., 1975; Peterson et al., 1988; Wheeler et al., 2003; Foster et al., 2009), Nickerson et al. (2002) captured 16 larval-sized (<130 mm) individuals out of 33 total Hellbenders in ~80 hours of survey effort in Little River, Tennessee. Over a ten-year period (2000-2010), the Hellbender population stage class structure in Little River remained consistent, with many larvae and relatively few large adults captured (Hecht-Kardasz et al., 2012). Crayfish, the principle component of the adult Hellbender diet (Nickerson and Mays, 1973a), also appeared to be scarcer and smaller in size in Little River than in other well-studied Hellbender localities (Nickerson et al., 2003). Nickerson et al. (2003) cited the low prey availability as a potential cause for the lack of large adult *C. alleganiensis* within Little River.

This study was conducted to determine if differences in prey availability could potentially impact the population structure of Hellbenders by influencing growth rates and potentially mortality rates. We assumed that if crayfish abundance impacts survival and/or growth of adult Hellbenders, a correlation would exist between body condition of adult Hellbenders and local crayfish abundance. Body condition was compared among three populations in streams with different crayfish abundance. We hypothesized that *C. alleganiensis* populations in localities with lower crayfish abundance would exhibit poorer body condition than individuals from rivers with higher crayfish abundance.

## Materials and Methods

### Study Sites

To evaluate the effect of prey abundance on Hellbender body condition, data from four studies completed in Little River since 2000 were compared to Nickerson and May’s (1973b) historic data from the North Fork of the White River in 1969, as well as data from 2004-2010 surveys of Hiwassee River, TN. These sites were chosen because of differences in prey availability based on preliminary data (Nickerson et al., 2003; Freake, unpubl. data).

The Little River study site was located entirely within the eastern Tennessee portion of Great Smoky Mountains National Park and is part of a regional reference water quality site. Lying within the Blue Ridge physiographic province, late Precambrian Elkmont and Thunderhead metamorphosed sandstone primarily comprised Little River’s bedrock (Mast and Turk, 1999). Large densities of dense rounded boulders, cobble, and gravel, formed from bedrock erosion, lie on the streambed. Interstitial habitat was limited within the streambed as sand filled in the majority of gravel beds. Elevation ranged from 327-407 m. Upland pine and river cove hardwood forest habitat was largely forested, with the exception of Scenic Highway TN 73, which ran adjacent to Little River, and several pull offs and parking lots. As part of the most visited national park in the country, Little River hosted heavy recreational use including fishing, swimming, kayaking, and inner tube users. During the summer of 2010, 281 tubers were counted within portions of the research site in five hours (Hecht, unpubl. data). Visitors also regularly disturbed stream substrate by building dams and channels and overturning rocks.

The streambed of the North Fork of the White River, a spring-fed river in Southern Missouri’s carbonate influenced Salem Plateau, was mostly a mixture of chert, dolomite, and sandstone (Nickerson et al., 2003). Limestone rock types are more susceptible to weathering and erosion than metamorphic rock and typically fragment and erode into the flat slabs. At the time of the study, little of the oak-hickory and oak-pine upland habitat in the research section had been cleared and siltation or embedding of the streambed particles was uncommon. Elevation ranged from ~198-202 m. During the study period, recreational human use such as swimming and canoeing was recorded but limited in scope.

The Hiwassee Scenic River study site was in the southeastern Tennessee portion of the Cherokee National Forest and was surrounded by mixed oak and oak-pine forest. This area is on the edge of the Blue Ridge Physiological Province and is largely comprised of clastic sedimentary rocks such as sandstone, siltstone, shale, and quartz-pebble conglomerate. Some lower reaches had limestone influence. Water flow in the study area was determined largely by releases from the Apalachia Dam which then flows in a 13 km tunneled flume to the Apalachia Powerhouse downstream. At full power, the powerhouse can release flows of cold hypolimnetic water reaching 80 m^3^/s. A minimal flow rate of 5.66 m^3^/s (200-cfs) was established in the 1990s. Water depths increased ~30-60 cm in some sites during high flow periods. The dam also influenced the streambed substrate. In sections heavily impacted by flow, rocks and particles were washed away, leaving only embedded rocks. Loose sediment gathered in some locations during low flow periods. *C. alleganiensis* was mostly limited to areas where the streambed was largely unaffected by flow rate. Due to the dam, the area attracts kayakers and white-water rafters. Fishing is also common in the area.

North Fork White River temperatures are similar to those in Little River for most of the summer and fall; winter and spring temperatures are often a few degrees above Little River. The North Fork White River site has large springs just upstream and within the research section, while large springs are not present in Little River. During the 2008 – 2010 study period, water temperature in Little River averaged 22.84 ± 2.03 °C (range: 14.60 – 22.80 °C; n = 103). During 1970 our year-round North Fork White River research sections water temperatures ranged from 9.8 °C (21 Feb) to 22.5 °C (8 July) N =104 (Nickerson and Mays 1973a; Nickerson, unpubl. data). A comparison of degree days on the North Fork White River watershed for five months 2004 – 2007 were warmer with earlier springtime temperatures than in 1969 – 1972 (Nickerson et al., 2009). The Hiwassee River research section is managed as a trout fishery (Young and Fiss 2005). The management plan is also known as “The 70 Degree Pledge”. Preferred temperature ranges for Rainbow Trout are 12 – 19 °C and Brown Trout 12 – 17 °C (Mettee et al., 1996). Tennessee Valley Authority took 320 grab water samples below the Appalachia Power House upstream from the Hiwassee research site between 4 July 1998 and 25 August 2017 (Baker 2017). Only seven samples were above 21.1 °C = 70 °F, and five of these were in October and taken at midday or afternoon. The highest was 23.81 °C on 26 October. Despite the demands for recreation and flood control, the Hiwassee section seems well managed for trout fishery temperatures. However, the systems flow is highly manipulated, and the temperature ranges may fluctuate greatly in a brief period.

### Field Methods

Diurnal skin diving was used to survey for Hellbenders in Little River in 2000 and 2004 – 2010, and Hiwassee River from 2004 – 2010. This method has been successful in locating all size classes of Hellbenders (Nickerson and Krysko, 2003). We turned rocks and other potential shelters, and captured Hellbenders by hand following detection. Surveys conducted by the Lee University research group used log peaveys in Hiwassee River and Little River to lift large boulders. Total length (TL) and snout-vent length (SVL) of each individual was measured in millimeters (mm). Mass was recorded in grams using an Ohaus® CS2000 compact digital scale (accuracy ±1.0 g; Ohaus Corporation, Parsippany, NJ, USA), DYMO® Pelouze SP5 digital scale (accuracy ±1.0 g; DYMO, Norwalk, CT, USA), or Pesola® spring scale (accuracy ±0.3%; Pesola AG, Baar, Switzerland). Sex was recorded if it could be determined based on presence or absence of swollen cloacal glands during August and September (Nickerson and Mays, 1973a). 9 mm or 12.5 mm Passive Integrated Transponder (PIT) tags (Destron-Fearing, South Saint Paul, MN, USA) were injected in adult and most sub-adult individuals dorsally near the base of the tail. Needles were sterilized with 70% ethanol between uses, and injection sites were treated with New Skin^®^ liquid bandage (Prestige Brands, Inc., Irvington, NY, USA).

Data from the North Fork of the White River used in this study was collected in 1969. Historical data was used for analysis because populations in this stream have declined drastically and are currently experiencing issues which could affect their growth and body condition (Nickerson et al., 2011; Bodinof et al., 2012). Methods utilized in the North Fork of the White River studies were similar to those conducted in Little River and Hiwassee River, with the exception that mammalian ear tags or Floy T-tags, rather than PIT tags, were used to individually mark animals, and mass was collected with an Ohaus triple beam scale (Ohaus Corporation, Parsippany, NJ. USA).

Prey abundance was measured by calculating crayfish relative frequencies as in Nickerson et al. (2003). Based on this method, the number of rocks turned were compared with those sheltering crayfish to calculate the percentage of rocks harboring crayfish. Crayfish relative frequency data were collected in 2009 for Little River, 2010 for the Hiwassee River, and during 1969 for the North Fork of the White River.

### Data Analysis

All data analysis was conducted in R (R Development Core Team, 2018). To evaluate the effect of prey abundance on Hellbender body condition data collected in Little River was compared to Nickerson and May’s (1973b) historic data from North Fork of the White River, and data collected from Hiwassee River. To linearize the relationship between TL and mass, we transformed all TL measurements by cubing and dividing by 10,000. To test differences in body condition among the three populations, we used analysis of covariance (ANCOVA) to compare linear regression models of transformed TL vs. mass of each Hellbender populations. The ANCOVA method was selected for use because it has received considerable attention and support in the scientific literature as a statistically accurate method for comparing body condition (see García-Berthou, 2001). Pair-wise tests were used to further examine body condition trends of the targeted Hellbender populations.

## Results

Total length of Hellbenders ranged from 40 – 520 mm in Little River, 40 – 460 mm in the Hiwassee, and 120 – 535 mm in the North Fork. Mean mass of all Little River Hellbenders (n=494) was 115.1g (±142.5) but was influenced by the large number of larval individuals (n=168). Mean mass of adults (n=183) was 266.6 g (±128.3). Hellbenders in the North Fork of the White River (n=463) had an average mass of 371.3 g (±240.4) (Nickerson and Mays, 1973b). Hiwassee Hellbenders (n=414) averaged 139.9 g (±123.6) in mass. Detailed accounts of population size structures can be found in Hecht-Kardasz et al. (2012) and Freake and DePerno, (2017).

Crayfish relative frequencies in Little River were low, ranging from 2-16% (mean=6.8% ± 3.8). At the North Fork of the White River site crayfish relative frequencies were considered high during the study period in 1969 and ranged from 56-67% (Nickerson and Mays, 1973a). Crayfish relative frequencies in Hiwassee River were intermediate and ranged from 21-28.5% with a mean of 24.7% ± 2.8.

Results of linear regression analysis of body condition in the three rivers are listed in Table 1. An ANCOVA comparing linear regression lines of body conditions in all three rivers (Fig. 1) was significant (F(2, 1490)=137.8, p<0.001). Individual pair-wise comparisons of Hellbender body condition in the three populations confirmed that the linear regression slope of Little River was significantly different from both North Fork of the White River (F(1,986)=194.6, p<0.001) and Hiwassee River (F(1, 1029)=16.6, p<0.001). The slope of Hellbender body condition in Hiwassee River was also significantly different from the slope of Hellbender body condition in North Fork of the White River (F(1,965)=92.5, p<0.001). Little River had the smallest expected mass per adjusted total length of the three rivers.

**Table 1.**
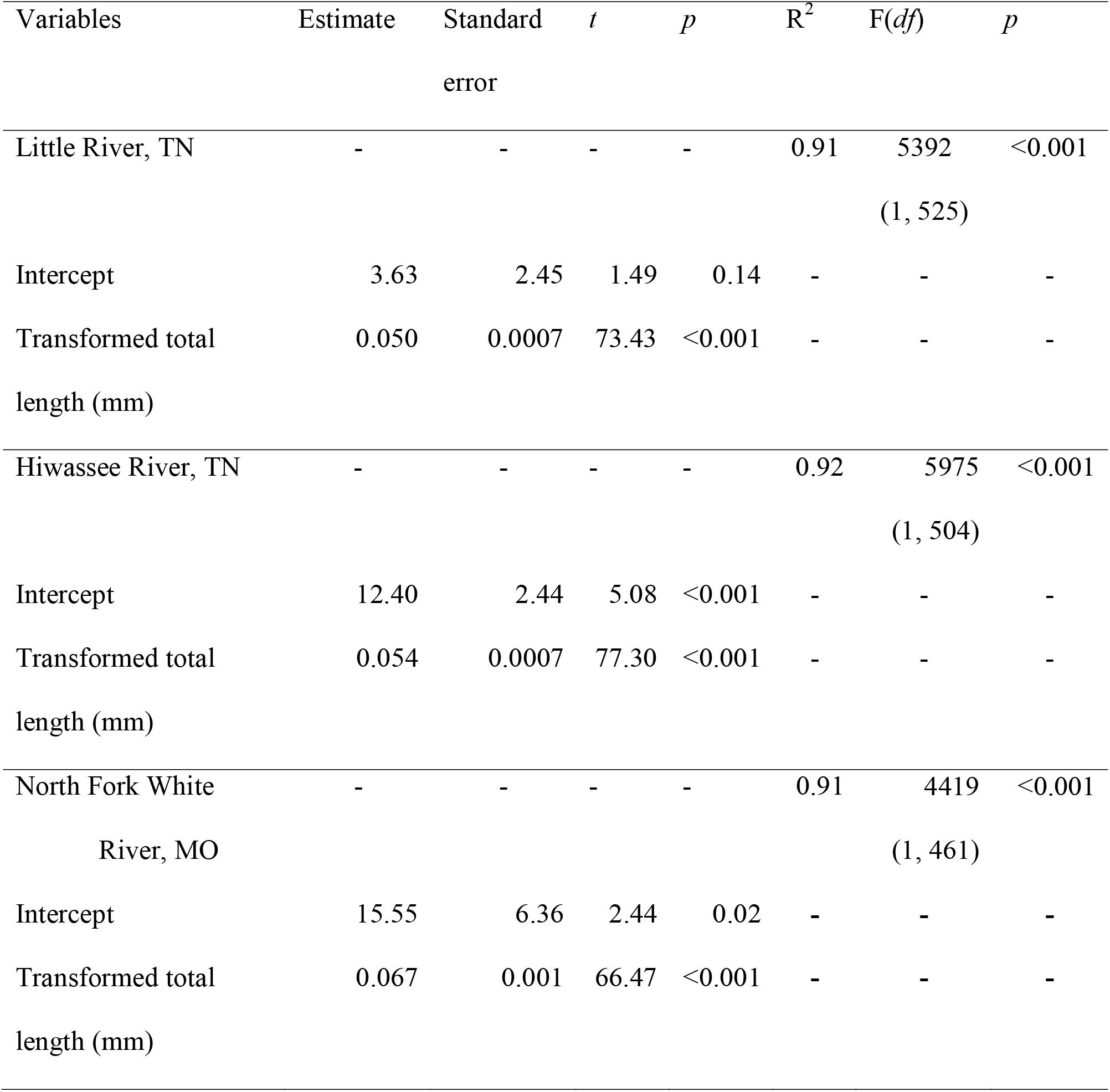
Variable estimates and model fit for linear regressions of Hellbender (*Cryptobranchus alleganiensis*) body condition (mass (g) vs. transformed total length (mm)) in three rivers.

**Fig 1.**
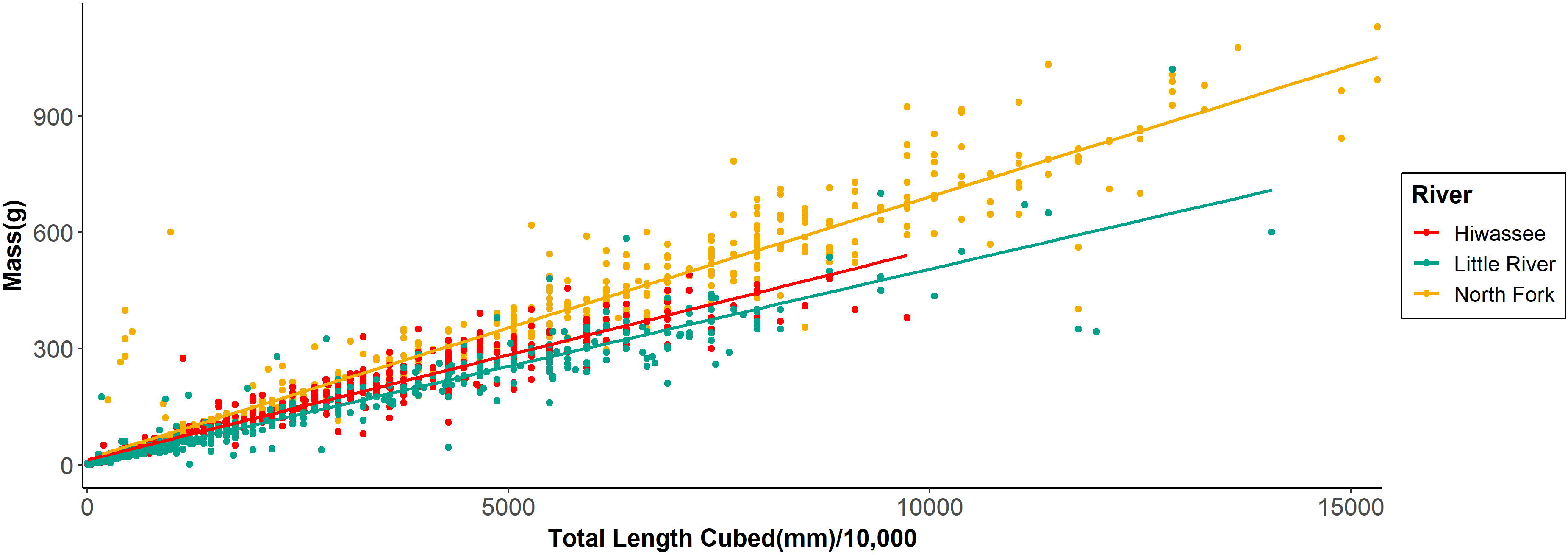
Scatter plot with regression lines comparing body condition of hellbenders (*Cryptobranchus alleganiensis*) from three rivers (Little River, Tennessee (n=527); Hiwassee River, Tennessee (n=507); North Fork of the White River; Missouri (n=463) with differing crayfish relative frequencies

## Discussion

The linear regression lines of Hellbender body condition in three rivers with different crayfish relative frequencies demonstrates that prey abundance correlates with overall body condition of *C. alleganiensis*. Crayfish relative frequency values appear to corroborate the observations of Bouchard (in Nickerson et al., 2003, p 623) correlating crayfish abundance with streambed rock composition. His observations suggested that rivers with non-carbonate bedrock typically exhibited lower crayfish densities. Crayfish relative frequencies in Little River, which has the least carbonate influence, were lower in comparison to the Hiwassee River and North Fork of the White River. While geology and system productivity may limit prey availability, it is important to note that Hellbenders may also influence top-down influence on prey abundance.

This study goes further to correlate the differences seen in prey abundance in these rivers to Hellbender body condition. Hellbenders in Little River showed a significantly lower linear regression slope when comparing trends in body condition between the three rivers, suggesting that Hellbenders in Little River are expected to have the lowest mass at a given total length of the three sites. North Fork of the White River, which is the only stream to have a major carbonate rock influence, had the highest crayfish relative frequencies as well as the highest overall Hellbender body condition. These trends were less clear in small individuals, which primarily feed on aquatic insect larvae rather than crayfish in Little River (Hecht et al., 2017). More recent studies in North Fork of the White River revealed an overall decrease in crayfish relative frequencies since 1969 (Nickerson et al., 2009), but no significant reduction in overall Hellbender body condition during the last 20 years (Wheeler et al., 2003). However, crayfish relative frequencies are still relatively large in comparison to both Little River and Hiwassee River.

Body condition can be affected by a host of factors including age, sex, parasites, infections, energy expenditures, behavioral changes, and various stressors, and ectotherms may experience quite different reactions to these factors than endotherms (Feder, 1991). Field conditions generally limit the focus of studies: mass and lengths of salamanders to two measures to assess body conditions: mass and lengths of salamanders taken from individuals of similar age and sex, collected at similar times and different sites to compare against temperatures and food sources. These data are still difficult to collect for Hellbenders.

In this study we compared data from the three study sites with food sources, but temperature, sex, age, and health are major considerations that could impact Hellbender body condition in our study areas. Temperatures for these rivers were similar in the three locations and do not explain the patterns seen in the body condition data. The highest and lowest average body condition were noted in two rivers with similar temperatures, while the stream with the coldest temperatures had the mid-range body condition. Sex and age could not be determined during the period, but a wide range of size classes were represented and the sex ratios of those able to be sexed were relatively equal in all three rivers (Nickerson and Mays, 1973a; Hecht-Kardasz, 2012; Freake and DePerno, 2017). For our study, health was not believed to play a large role in Hellbender body conditions within the rivers as little evidence for major health problems was present in these populations at the time of data collection. Similarly, injury did not seem to be a significant issue in the study populations when the data were collected. In 1969 only 2.9% of 479 individuals in North Fork White River were abnormal/injured and they exhibited rapid regeneration (Hiler et al., 2005) and additional surveys between 1972 and 1980 continued to show immense and healthy populations (Nickerson and Briggler, 2007). Hellbenders in Little River and Hiwassee River appeared in good overall health with few major injuries or external signs of disease. Common abnormalities were minor and included missing digits, extra digits and scars. Only a very small number of potentially serious abnormalities such as ulcers and severe limb injuries were noted. A study in Little River by Souza et al. (2012) found Bd and Ranavirus DNA was detected in 31% and 19% of tested Hellbenders respectively, although noted abnormalities were not consistent with chytridiomycosis or ranaviral disease.

While we cannot conclusively rule out the influence of other factors, this study provides support that prey abundance may be a limiting factor for Hellbender body condition in rivers with low crayfish densities. Adult Hellbenders in Little River often had empty stomachs when palpated during the summer months (Hecht, pers. obs.). Additional study into adult Hellbender diet in Little River may provide additional insight on whether individuals switch their prey to consume other, potentially less nutritious organisms as the main portion of their diet in Little River or if they are simply eating less. Currently, Hellbenders do not appear unhealthy in the protected portions of Little River and are reproducing regularly. However, the strain of low body condition could become a larger conservation concern if the population faces additional stressors or loses additional prey base in the future so further study and periodic monitoring of the health and status of the Hellbender population there should continue.

## Acknowledgments

We would like to thank M. Souza, The Great Smoky Mountains Institute at Tremont, P. Ross, M. Christman, K. Dodd, the Williams Lab at Purdue University, and all volunteers for assistance on this project. We would also like to acknowledge P. Super, K. Langdon, and the National Park Service. T. Baker and the Tennessee Valley Authority provided temperature information for the Hiwassee River. Lastly, we would like to thank Gina Hillsberry for editing and formatting assistance on the manuscript. Financial support for this research was provided by the Great Smoky Mountains Conservation Association: Carlos C. Campbell Fellowship, The Reptile and Amphibian Conservation Corp (RACC), the Cryptobranchid Interest Group: Jennifer Elwood Conservation Grant, U.S. Geological Survey (U.S. Department of Interior), Max Allen’s Zoological Gardens, and the Arkansas State Faculty Grant. Research was conducted under permits from the National Park Service (GRSM-2009-SCI-0061, GRSM-2009-0056, GRSM-2008-SCI-0052, GRSM-00-0131, GRSM-2004-0307), and scientific collecting permits from the Missouri Department of Conservation. Study was conducted with the approval of the Institutional Review Board of Lee University, University of Florida IACUC (#A560), and University of Florida ARC Protocol (#017-08WEC).

## Literature Cited

Anderson, R. O, and S. J. Gutreuter. 1983. Length, weight, and associated structural indices. Pp. 283 – 300 in L.A. Nielson, and D.L. Johnson (Eds.), Fisheries Techniques. American Fisheries Society, Maryland, USA.

Bodinof, C. M., J. T. Briggler, R. E. Junge, T. Mong, J. Beringer, M. D. Wanner, C. D. Schutte, J. Etting, and J. J. Millspaugh. 2012. Survival and body condition of captive-reared juvenile Ozark Hellbenders (*Cryptobranchus alleganiensis bishopi*) following translocation in the wild. Copeia 2012:150 – 159.

Davis, A. K., and J. C. Maerz. 2007. Spot symmetry predicts body condition in spotted salamanders, *Ambystoma maculatum*. Applied Herpetology 4:195 – 205.

Feder, M. E. 1991. A perspective on environmental physiology of the amphibians. In. Environmental Physiology of the Amphibians. 1 – 6 p. University of Chicago Press, viii + 1 - 646.

Freake, M. J., and C. S. DePerno. 2017. Importance of demographic surveys and public lands for the conservation of eastern hellbenders *Cryptobranchus alleganiensis* in southeast USA.PLoS ONE 12(6): e0179153. https://doi:org/10.1371/journal.pone.o179153

Foster, R. L., A. M. Mcmillan, and K. J. Roblee. 2009. Population status of Hellbender salamanders (*Cryptobranchus alleganiensis*) in the Alleghany River Drainage of New York State. Journal of Herpetology 43:579 – 588.

García-Berthou, E. 2001. On the misuse of residuals in ecology: testing regression residuals vs. the analysis of covariance. Journal of Animal Ecology 70:708 – 711.

Gendron, A. D., D. J. Marcogliese, S. Barbeau, M. S. Christin, P. Brousseau, S. Ruby, D. Cyr, and M. Fournier. 2003. Exposure of leopard frogs to a pesticide mixture affects life history characteristics of the lungworm *Rhabdias ranae*. Oecologia, 135(3):469 – 476.

Girish, S., and S. K. Saidapur. 2000. Interrelationship between food availability, fat body, and ovarian cycles in the frog, *Rana tigrina*, with a discussion on the role of fat body in anuran reproduction. Journal of Experimental Zoology 286:487 – 493.

Hecht, K. A., M. A. Nickerson, and P. B. Colclough. 2017. Hellbenders (*Cryptobranchus alleganiensis*) may exhibit an ontogenetic dietary shift. Southeastern Naturalist 16(2):157 – 162.

Hecht-Kardasz, K. A, M. A. Nickerson, M. Freake, and P. Colclough. 2012. Population structure of the Hellbender (*Cryptobranchus alleganiensis*) in a Great Smoky Mountains stream. Bulletin of the Florida Museum of Natural History 51:227 – 241.

Hiler, W. R., B. A. Wheeler, and S. E. Trauth. 2005. Abnormalities in the Ozark Hellbender (*Cryptobranchus alleganiensis bishopi*) in Arkansas: A comparison between two rivers with a historical perspective. Journal of Arkansas Academy of Science 59:88 – 94.

Homyack, J. A., C. A. Haas, and W. A. Hopkins. 2011. Energetics of surface□active terrestrial salamanders in experimentally harvested forest. The Journal of Wildlife Management 75(6):1267 – 1278.

Howard, R. D., and J. R. Young. 1998. Individual variation in male vocal traits and female mating preferences in *Bufo americanus*. Animal Behavior 55;1165 – 1179.

Howard, R. D., R. S. Mooyman, and H. H. Whiteman. 1997. Differential effects of mate competition and mate choice on eastern tiger salamanders. Animal Behavior 53: 1345 –1356.

Humphries, W. J, and T. K. Pauley. 2005. Life history of the Hellbender, *Cryptobranchus alleganiensis*, in a West Virginia stream. American Midland Naturalist 154:125 – 142

Karraker, N. E., and H. H. Welsh Jr. 2006. Long-term impacts of even-aged timber management on abundance and body condition of terrestrial amphibians in Northwestern California. Biological Conservation 131(1):132 – 140.

Levey, R., N. Shambaugh, D. Fort, and J. Andrews. 2003. Investigations into the causes of amphibian malformations in the Lake Champlain Basin of New England. Waterbury, VT: Vermont Department of Environmental Conservation.

Lowe, W. H., G. E. Likens, and B. J. Cosentino. 2006. Self-organisation in streams: the relationship between movement behavior and body condition in a headwater salamander. Freshwater Biology 51:2052 – 2062.

Mast, M. A., and J. T. Turk. 1999. Environmental characteristics and water quality of Hydrologic Benchmark Network stations in the Eastern United States, 1963-95: U.S. Geological Survey Circular 1173-A, 158 p.

Mettee, M. F. P. E, O’Neil, and J. M. Pierson. 1996. Fishes of Alabama and the Mobile Basin. State of Alabama, Oxmoor House, Inc.

Nickerson, M. A., and C. E. Mays. 1973a. The Hellbenders: North American Giant Salamanders. Milwaukee Public Museum, Milwaukee, WI, 106 p.

Nickerson, M. A., and C. E. Mays. 1973b. A study of the Ozark Hellbender, *Cryptobranchus alleganiensis bishopi*. Ecology 54:1164 – 1165.

Nickerson, M. A., and K. L. Krysko. 2003. Surveying for Hellbender salamanders, *Cryptobranchus alleganiensis* (Daudin): A review and critique. Applied Herpetology 1: 37 – 44.

Nickerson, M. A., and J. T. Briggler. 2007. Harvesting as a factor in population decline of a long-lived salamander; the Ozark Hellbender, *Cryptobranchus alleganiensis* Grobman. Applied Herpetology 4:207 – 216.

Nickerson, M. A., K. L. Krysko, and R. D. Owen. 2002. Ecological status of the Hellbender (*Cryptobranchus alleganiensis*) and the mudpuppy (*Necturus maculosus*) salamanders in Great Smoky Mountains National Park. Journal of the North Carolina Academy of Science 118:27 – 34.

Nickerson, M. A., K. L. Krysko, and R. D. Owen. 2003. Habitat differences affecting age class distributions of the Hellbender salamander, *Cryptobranchus alleganiesnsis*. Southeastern Naturalist 2:619 – 629.

Nickerson, M. A., A. L. Pitt, and J. T. Tavano. 2009. Decline of the Ozark Hellbender (*Cryptobranchus alleganiensis bishopi*) in the North Fork of White River, Ozark County, Missouri: A historical habitat perspective. A final report submitted to the Saint Louis Zoological Park and the Reptile and Amphibian Conservation Corp. Reptile and Amphibian Conservation Corps: Gainesville, FL, 53 p.

Nickerson, C. A., C. M. Ott, S. L. Castro, V. M. Garcia, J. Briggler, A. L. Pitt, J. K. Byram, and M. A. Nickerson. 2011. Evaluation of microorganisms cultured from injured and repressed tissue regeneration sites in endangered giant aquatic Ozark Hellbenders salamanders. PLoS ONE 6(12):e28905.

Peterson, C. L., D. E. Metter, B. T. Miller, R. F. Wilkinson, and M. S. Topping. 1988. Demography of the Hellbender *Cryptobranchus alleganiensis* in the Ozarks. American Midland Naturalist 119:291 – 303.

Pope, K. L., and K. R. Matthews. 2002. Influence of anuran prey on the condition and distribution of *Rana muscosa* in the Sierra Nevada. Herpetologica 58:354 – 363.

R Development Core Team. 2018. R: A language and environment for statistical computing. R Foundation for Statistical Computing, Vienna, Austria. Retrieved from http://www.R-project.org.

Reading, C. J, and R. T. Clarke. 1995. The effects of density, rainfall, and environmental temperature on body condition and fecundity in the Common Toad, *Bufo bufo*. Oecologia 102:453 – 459.

Souza, M. J., M. J. Gray, P. Coclough, and D. L. Miller. 2012. Prevalence of infection by *Batrachochytrium dendrobatidis* and *Ranavirus* in eastern Hellbenders (*Cryptobranchus alleganiensis alleganiensis*) in eastern Tennessee. Journal of Wildlife Diseases 48:560 – 566.

Stevenson, R. D., and W. A. Woods, Jr. 2006. Condition indices for conservation: New uses for evolving tools. Integrative and Comparative Biology 46:1169 – 1190.

Taber, W. E., R. F. Wilkinson, Jr., and M. S. Topping. 1975. Age and growth of Hellbenders in the Niangua River, Missouri. Copeia 1975:633 – 639.

Welsh, Jr., H. H., K. L. Pope, and C. A. Wheeler. 2008. Using multiple metrics to assess the effects of forest succession on population status: a comparative study of two terrestrial salamanders in the US Pacific Northwest. Biological Conservation 141:1149 – 1160.

Wheeler, B. A., E. Prosen, A. Mathis, and R. F. Wilkinson. 2003. Population declines of a long-lived salamander: A 20+ year study of Hellbenders, *Cryptobranchus alleganiensis*. Biological Conservation 109:151 – 156.

Young, D., and F. Fies. 2005. Management Plan for the Hiwassee River Trout Fishery. 2005 – 2010. Tennessee Wildlife Resources Agency 19 p.

